# Charge-voltage curves of Shaker potassium channel are not hysteretic at steady state

**DOI:** 10.1101/2022.11.07.515208

**Authors:** John Cowgill, Baron Chanda

## Abstract

Charged residues in transmembrane segments of voltage-gated ion channels (VGICs) sense and respond to changes in the electric field. The movement of these gating charges underpins voltage-dependent activation and is also a direct metric of the net free-energy of channel activation. However, for most voltage-gated ion channels, the charge-voltage (Q-V) curves appear to be dependent on initial conditions. For instance, Q-V curves of Shaker potassium channel obtained by hyperpolarizing from 0 mV is left shifted compared to those obtained by depolarizing from a holding potential of -80 mV. This hysteresis in Q-V curves is a common feature of channels in the VGIC superfamily and raises profound questions about channel energetics because the net free-energy of channel gating is a state function and should be path independent. Due to technical limitations, conventional gating current protocols are limited to test pulse durations of ≤500 ms which raises the possibility that the dependence of Q-V on initial conditions reflects a lack of equilibration. Others have suggested that the hysteresis is fundamental thermodynamic property of voltage-gated ion channels and reflects energy dissipation due to measurements under non-equilibrium conditions inherent to rapid voltage jumps (Villalba-Galea, 2017). Using an improved gating current and voltage-clamp fluorometry protocols, we show that the gating hysteresis arising from different initial conditions in Shaker potassium channel is eliminated with ultra-long (18-25 second) test pulses. Our study identifies a modified gating current recording protocol to obtain steady state Q-V curves of a voltage-gated ion channel. Above all, these findings demonstrate that the gating hysteresis in Shaker channel is a kinetic phenomenon rather than a true thermodynamic property of the channel and the charge-voltage curve is a true measure of the net-free energy of channel gating.

## Introduction

Ion channels regulate electrical signaling by gating in response to changes in mechanical pressure, temperature, noxious chemicals, secondary messengers, and/or membrane potential. From a thermodynamic standpoint, ion channels are molecular machines that harness various forms of energy to drive a conformational change that results in opening or closing of an ion permeation pathway through the cell membrane. Therefore, at equilibrium, the free energy difference between the initial (resting) and final (fully activated) state can be determined by measuring the reversible work done on the system (Chowdhury and Chanda, 2012; Sigg, 2013). Analogous to mechanical work, the net free energy change in the system in response to a stimulus can be calculated by measuring the conjugate displacements corresponding to the thermodynamic potential (Table 1). For a ligand-dependent signaling pathway, the work function associated with ligand-dependent gating can be estimated from ligand binding curves (Wyman and Gill, 1990). The ligand binding curve is identical to the dose-response curves for channels if and only if the channel gating can be described using a two-state model. We should note that for most ion channels dose-response curves do not overlap with binding curves indicating that channel gating involves additional states (Auerbach, 2012; Changeux and Christopoulos, 2016; Changeux et al., 1984; Horrigan and Aldrich 2002). In such instances, the free energy of channel gating cannot be estimated by using the dose-response curves (Chowdhury and Chanda, 2012; Sigg, 2013).

**Table 1:** List of forces and conjugate displacements for different channel types

| Type of Channel | Force | Displacement |
| --- | --- | --- |
| Ligand-gated | Chemical potential ( $\mu$ ) | Fraction bound ( $X$ ) |
| Mechanosensitive |  |  |
| Volume-regulated | Pressure ( $P$ ) | Volume ( $V$ ) |
| Stretch-activated | Surface tension ( $\gamma$ ) | Area ( $A$ ) |
| Elastic | Force ( $f$ ) | Length ( $L$ ) |
| Thermosensitive | Temperature ( $T$ ) | Entropy ( $S$ ) |
| Voltage-gated | Voltage ( $V$ ) | Charge ( $Q$ ) |

For a voltage-gated ion channel, the conjugate pairs for calculating the work function are the applied voltage and gating charge movement (Table 1). Therefore, gating charge vs. voltage curves (Q-V) provide a direct metric of the net free energy of channel activation (Chowdhury and Chanda, 2012) (Sigg, 2013). A fundamental property of free-energy is that it is a state-function and, hence, the free-energy difference is only determined by the difference between the two states and is independent of the path. Therefore, if the charge voltage curves are a measure of the free energy difference between the resting and activated state, then the Q-V curves measured from different holding potentials should be identical at equilibrium. Experimentally measured Q-V curves, however, are profoundly influenced by starting conditions (Armstrong and Bezanilla, 1977; Bezanilla et al., 1991; Haddad and Blunck, 2011; Lacroix et al., 2011; Olcese et al., 1997; Shi et al., 2019; Villalba-Galea, 2017; Villalba-Galea et al., 2008). For instance, Q-V curves obtained by depolarizing test pulses for the Shaker potassium channel are right shifted compared to those obtained by hyperpolarizing pulses ((Bezanilla et al., 1982; Olcese et al., 1997) and Figure 1). This phenomenon, referred to as gating hysteresis, is a hallmark of nearly all voltage-gated ion channels and has also been observed in mechanosensitive (Nakayama et al., 2018; Yang et al., 2013), thermosensitive (Liu et al., 2011; Sánchez-Moreno et al., 2018), and ligand-gated channels (Brookes, 1980; Chang et al., 1984; Tilegenova et al., 2017).

**Figure 1:**
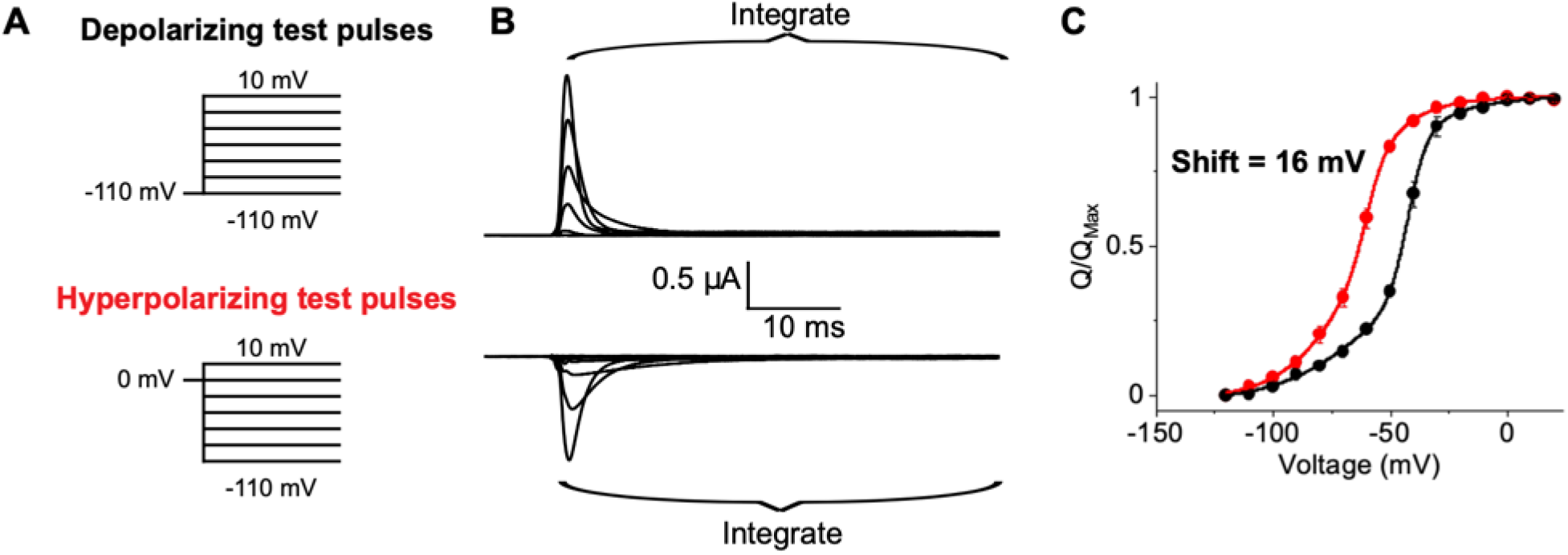
Hysteresis in traditional gating current recordings. (A) Traditional pulse protocols used to measure gating currents from channels. (B) Capacitive- and leak-subtracted gating current recordings for Shaker W434F using 50 ms test pulses measured in the depolarizing (top) or hyperpolarizing (bottom) direction. Both recordings are from the same oocyte and shown on the same scale for direct comparison. In the polarized holding potential condition, a 200 ms prepulse to -110 mV from a holding potential of -80 mV was used to minimize the time the oocyte was held at extreme potentials. The region integrated to quantify gating charge is indicated by the brackets. (C) Charge-voltage (Q-V) relationships for Shaker W434F recorded with variable test pulse durations with depolarizing (black) or hyperpolarizing (red) test pulses. The shift reported in each graph represents the difference in the median voltage extracted between the two curves. Error bars show standard error from 3 oocytes.

Hysteresis in Q-V curves (or any force-displacement curve) is confounding as it implies that the energies of channel opening and closing are different. When it was first observed, Bezanilla *et al*. (1982) speculated that “If the integration period were extended indefinitely, the Q-V curve would correspond to the steady-state distribution of charge vs. holding potential and it would not depend on the initial conditions”. This holding potential dependent shift in the Q-V curves is generally attributed to slow, especially voltage-independent processes that do not equilibrate within the timescale of traditional gating current recordings (Haddad and Blunck, 2011; Shi et. al. 2019; Lacroix et. al., 2011). Recently however, Villalba-Galea (2017) argued that hysteresis in charge voltage curves is an inherent property of the system and reflects a loss of energy during the activation process.

Thermodynamics dictates that the free-energy change in the system equals to the reversible work only and only when the process is carried out infinitesimally slowly. Since this requirement is never satisfied in practice, the observed hysteresis in the charge voltage curves must reflect a loss of energy due to the irreversible nature of work performed on the system. Without going into the merits of this argument, it is evident that such reasoning has the potential to invalidate the use of standard thermodynamic and kinetic analysis of channel gating. Principle of microscopic reversibility dictates that the free-energy required for activation is identical to the free-energy required for deactivation. We note that using any of the established kinetic models for Shaker channel gating (Zagotta et. al. JGP (1994), Bezanilla et. al. Biophys J, Schoppa and Sigworth JGP (1998)) it is not possible to simulate hysteresis in steady state Q-V curves. No energy is lost or gained when the channel goes through a gating cycle in any of these models. It may be possible to introduce hysteresis by arbitrarily truncating the integration times for simulated gating currents or by introducing asymmetric noise in the simulations but these do not change the fact that predicted steady state Q-V curves inherently do not exhibit any gating hysteresis.

However, the possibility that the channel gating is coupled to an additional energy-generating pathway cannot be ruled out simply based on theoretical considerations. All Q-V curves are obtained in living cells and there are multitude of processes, many of which can potentially provide energy for channel gating. Although channel activity has been measured in reconstituted systems, we are not aware of careful thermodynamic studies to rule the contribution of alternate pathways. Here, we reexamine the issue of gating current hysteresis by testing the proposition that the hysteresis simply reflects the inability of standard protocols to measure all the gating charge movement. If the charge movement is slow, then the gating current signal is small since gating current is first derivative of charge movement. As a result, it is difficult to resolve gating current signal from baseline noise and endogenous leak currents by simply extending the pulse duration. These technical considerations have historically placed an upper bound to the pulse length duration of around 500 ms for gating current recordings. Using a redesigned gating current recording protocol, we are able to measure slow charge movements some of which take tens of seconds to complete. We demonstrate that these steady state charge-voltage (Q_ss_-V) measurements are independent of the initial holding potential as expected for a thermodynamic state function. Furthermore, by spectroscopically tracking the voltage sensor movement using voltage-clamp fluorometry (VCF) under conditions that minimize photobleaching, we also show that the F-V curves in forward and reverse direction are superimposable. Together, these findings support the notion that Q-V curve obtained under equilibrium conditions is a true measure of the net free-energy of voltage-activation and that the established gating models are sufficient to fully describe the thermodynamic properties of Shaker potassium channel.

## Methods

All mutations were introduced to the fast inactivation-removed Shaker potassium channel (Δ6–46) using QuikChange mutagenesis. The W434F mutation was inserted into all constructs to render the channel nonconductive while the A359C mutation was added to the construct used in VCF studies to enable fluorophore labeling. All constructs were confirmed across the entire open reading frame by Sanger sequencing. Constructs were linearized using NotI digestion (Fermentas-Thermo Fisher Scientific) and transcribed *in vitro* with the mMESSAGE mMACHINE T7 kit (Life Technologies).

Stage IV oocytes were surgically removed from *Xenopus laevis* following protocols approved by the University of Wisconsin-Madison Institutional Animal Care and Use Committee. Follicular layer was removed by treatment with 0.8 mg/mL collagenase (Roche) for 1–1.5□hours at room temperature. Oocytes were injected with 30□nl cRNA at a concentration of 1 μg/μl and incubated in ND-96 solution supplemented with 100 U/ml penicillin-streptomycin, 50□mg/ml tetracycline, 0.1□mg/ml amikacin, 50 μg/ml ciprofloxacin, 100 μg/ml gentamicin and 0.5 mg/mL BSA at 18 °C before the recording. Gating current and VCF measurements were performed 2–4□days after injection. For VCF recordings, oocytes were labeled with 10 μM tetramethylrhodamine (TMR) maleimide (TMRM; Invitrogen) in a depolarizing solution (110 mM KCl, 1.5 mM MgCl_2_, 0.8 mM CaCl_2_, and 10 mM HEPES, pH 7.5) at 18°C for 30 minutes prior to use.

### Electrophysiology

All recordings were performed on a Dagan CA-1B amplifier in either cut-open voltage clamp (COVG) mode for gating current recordings or two-electrode voltage clamp (TEVC) mode for VCF recordings as previously described (Fernandez-Marino et al., 2018). Permeant ion-free external solution (105□mM NMG, 10□mM HEPES, 2□mM CaOH, pH 7.4) was used for all recordings. For recordings in COVG mode, the internal solution was 105□mM NMG, 10□mM HEPES, 5□mM EGTA, pH 7.4 and oocytes were permeabilized using internal solution supplanted with 0.3% saponin. The recording pipette was filled with 3□M KCl and broken to resistance of 200 to 800 kΩ prior to oocyte impalement. For gating current recordings, analog signals were sampled at 50□kHz with using a Digidata 1440□A A/D converter (Axon Instruments) and low-pass filtered at 10□KHz. VCF signals were recorded at a sampling frequency of 10 kHz. Holding potential and test pulse duration varied according to the descriptions in the figure legends. Interpulse durations of 4 seconds up to 2 minutes were used depending on the duration of the test pulse. All oocytes were measured at both depolarized and hyperpolarized holding potentials for a single test pulse duration to generate paired recordings. The Q-V curves were normalized to the maximum gating charge observed in a given protocol and the difference between the Qmax obtained with different holding potentials was negligible (see attached Figure S5).

### Voltage clamp fluorometry

Excitation was provided by an LED source (X-Cite) operated at minimum (5%) output at 505-545 nm through a 20X/0.46NA objective (Olympus). The illumination intensity was further reduced using a 0.4 ND (Semrock) filter to minimize photobleaching and filtered through a 525/50 nm bandpass filter (Semrock). A 562 nm dichroic (Semrock) and a 593/46 nm bandpass filter (Semrock) were used to separate fluorescence emission from reflected light. Fluorescence signals were detected with a Hamamatsu photomultiplier tube (PMT) detector and transmitted directly to the Digidata 1440□A A/D converter at a frequency of 10 kHz. Oocytes were recorded from a 0 mV holding potential, or -60 mV holding potential with a -110 mV prepulse for 500 ms in paired recordings as was done for gating current recordings. Interpulse duration was 2 minutes and test pulse duration was 25 seconds for all recordings.

### Data processing and statistical analysis

For gating current recordings, leak current was subtracted by adjusting the current at the end of the tail pulse to zero. Following baseline subtraction, the first 50 ms of the tail pulse was integrated using the built-in integration function in Clampfit (Molecular devices). Capacitive charge was removed following integration by subtraction of the slope of the charge-voltage relationship at depolarized potentials (Aggarwal and MacKinnon, 1996). VCF recordings were baseline subtracted prior the test pulse to remove the interpulse drift and then normalized to the average fluorescence value at the end of the -110 mV test pulse. Following normalization of the Q-V and F-V curves, median voltage was extracted as previously described using pseudo-integration via trapezoidal approximation (Chowdhury and Chanda, 2012). Free energy values were computed from median voltages extracted from Q-V curves as described previously assuming a Q_Max_ of 13 e^-^ (Aggarwal and MacKinnon, 1996). Statistical significance in the differences between median voltages was performed using the built-in paired T-test analysis in Excel 2016 using a significance cutoff of p<0.05. The number of independent observations represents the number of oocytes measured under paired conditions (fixed test pulse duration with depolarized and hyperpolarized holding potentials).

For the example traces in Fig. 2, capacitive current was subtracted from the recording offline. This was accomplished by fitting a capacitive transient from a +10 mV pulse from 0 mV to a modified Gaussian in Origin and scaling accordingly to subtract from all test pulses.

**Figure 2:**
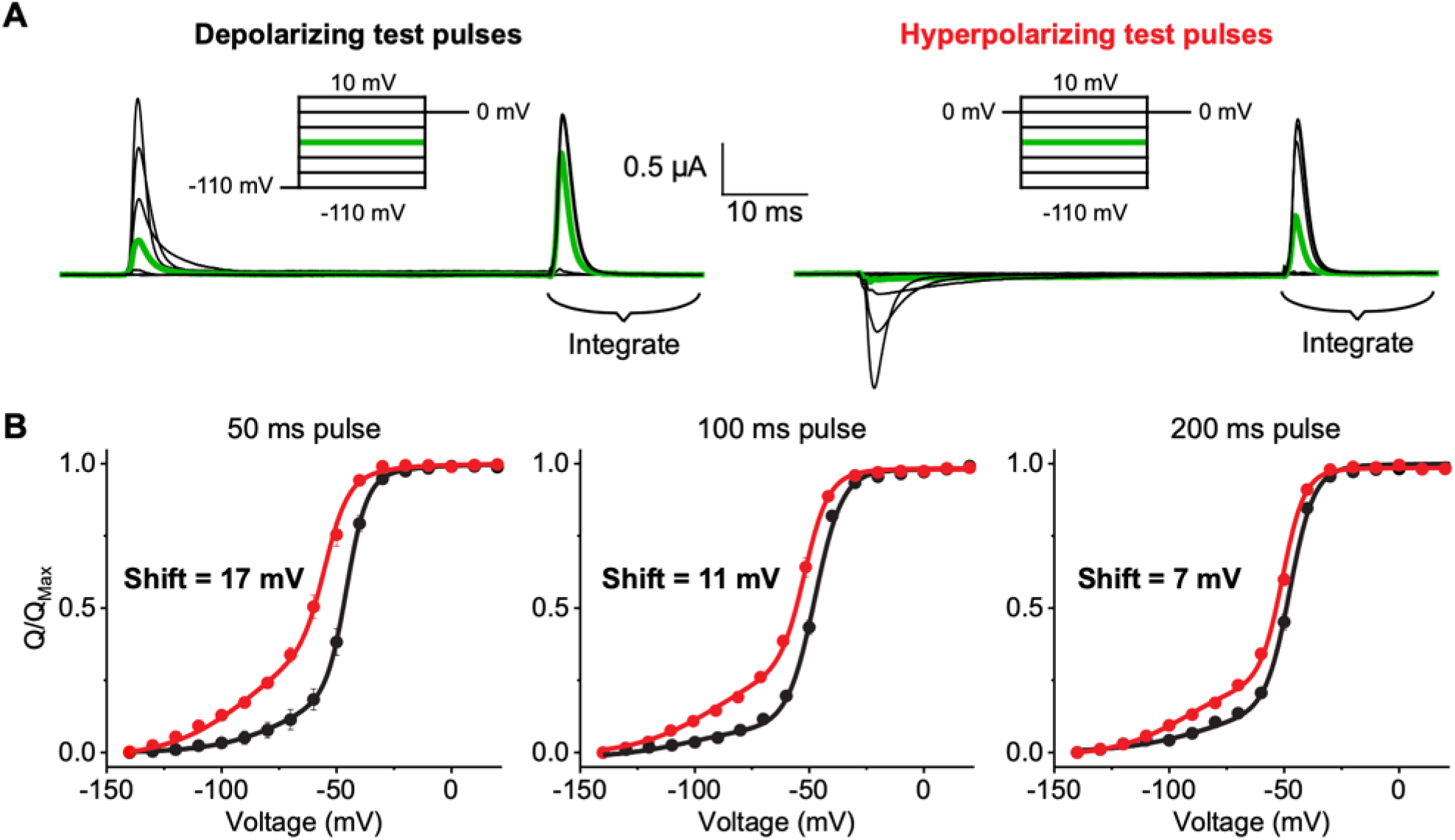
Hysteresis in Shaker gating decreases with increasing pulse duration. (A) Capacitive- and leak-subtracted gating current recordings using 50 ms test pulses from polarized holding potentials (left) or depolarized holding potentials (right). Both recordings are from the same oocyte and shown on the same scale for direct comparison. The off-gating pulse was used for integration of gating charge and was set to 0 mV due to fast gating current kinetics and minimal ionic current contamination. In the polarized holding potential condition, a 200 ms prepulse to -110 mV from a holding potential of -80 mV was used to minimize the time the oocyte was held at extreme potentials. (B) Charge-voltage (Q-V) relationships for Shaker W434F recorded with variable pulse durations in depolarizing (black) or hyperpolarizing (red) direction. The shift reported in each graph represents the difference in the median voltage extracted from the polarized and depolarized conditions. Error bars show standard error for at least 3 oocytes for each condition.

Kinetic model simulations were performed using the software Kinetic Model Builder version 2.0 (Goldschen-Ohm et al., 2014). Briefly, occupancy of each state was determined by numerical solution of the transition matrices within the model as described previously (Colquhoun and Hawkes, 1995).

## Results

### Improved protocol for gating current recordings

In a typical Q-V curve measurements (Fig. 1 and (Armstrong and Bezanilla, 1977)), the membrane potential is stepped from a single holding potential to a series of test potentials and gating charge is measured by integration of the current elicited during the test pulse. This gating charge represents the difference in charge state of a channel at all test potentials relative to a single reference point at the holding potential. While this represents the most common method for obtaining Q-V curves, this approach has drawbacks that can potentially complicate estimation of the charge moved. For instance, at potentials where kinetics of charge movement is slow, the current amplitude is small and some of the charge movement is eventually lost in the baseline (Fig. 2 A). In addition, the currents are being integrated at many different voltages, leading to variable amounts of contamination from ionic currents and drifts in the membrane/seal resistance. These errors are further compounded when the pulse length duration is increased.

An alternative strategy is to integrate the OFF gating currents upon return to the holding potential to estimate the amount of charge moved at various test potentials (Armstrong and Bezanilla 1974). Conservation of charge stipulates that this is equal in magnitude to the gating charge that was moved during the test pulse (Bezanilla, 2018). As the gating charge is measured at a single potential for all test pulses, this protocol reduces the contamination from non-linear current that plagues conventional gating current recordings (Goodchild and Fedida, 2014; Piper et al., 2003; Thouta et al., 2017). This strategy has been successfully applied to slowly activating channels like hERG (Shi et al., 2019; Thouta et al., 2017) as well as BK channels (Horrigan and Aldrich 2002) to expand the timescale of gating current recordings.

The conventional choice for a reference state has been to return to the holding potential following the test pulse. However, a better strategy is to return to a potential that maximizes the speed of charge movement while minimizing contaminating signals. Kinetics of gating charge movement depends exponentially on voltage, and can be rapidly accelerated in the extreme depolarizing or hyperpolarizing regions of the Q-V curve. However, endogenous currents and membrane instability give rise to contaminating signals outside the range of approximately -100 mV to 40 mV, For Shaker, we used 0 mV as the gating charge movement is nearly complete within 10 ms (Figure 2A) and the pseudo-symmetric solutions used minimize any ionic current contamination.

Furthermore, the normalized gating charge measured at 0 mV is the same for a 50 ms integration window (i.e. the points on the Q-V curves superimpose) whether depolarizing or hyperpolarizing pulses are used (Figure 1C). Thus, a 50 ms integration window approximates equilibrium at 0 mV. In addition to increasing signal to noise in gating current recordings, this protocol minimizes the timescale needed for gating current integration thereby limiting artifacts that typically contaminate such recordings. This protocol for measuring gating currents will henceforth be referred to as return to reference protocol.

The advantage of this protocol is clear in the test pulse to -50 mV from a holding potential of 0 mV in the example recording of Fig. 2 A (green trace on right). During the test pulse there is very little displacement from baseline, giving the appearance that no gating charge moves between 0 mV and -50 mV. However, upon return to 0 mV at the end of the pulse, a large gating current is observed. All of this charge moved during the 50 ms test pulse but was so diffuse that it results in a barely discernable gating current.

### Gating charge movement probed with variable test pulse duration

We first recorded gating currents from oocytes expressing Shaker W434F (Perozo et al., 1993) using the return to reference protocols (Fig 2 B). The two Q-V curves obtained by depolarizing or hyperpolarizing test pulses in response to a 50 ms pulse are displaced relative to each other as has been reported previously using standard pulse protocols (Lacroix et al., 2011). Furthermore, the magnitude of separation between median voltages (V_M_) of the curves decreases with increasing test pulse duration, ranging from 17 mV at 50 ms to 7 mV at 200 ms. This difference in V_M_ corresponds to an apparent free energy difference of 5.2 kcal/mole for a 50 ms pulse and 2 kcal/mole for a 200 ms pulse. Considering the net activation energy of Shaker W434F is only -14 kcal/mole (Chowdhury and Chanda, 2012), these differences in measured free energies are substantial.

If the difference in Q-V curve using standard protocol (Fig. 1C) reflects additional stabilization of the voltage-sensor in the up conformation due to voltage sensor relaxation or any such voltage-independent epiphenomenon, it is unclear why its magnitude is dependent on pulse length. In principle, any transition that specifically stabilizes either the resting or activated state of the channel should be reflected in the measured Q-V curves (see Appendix). Next, we consider the possibility that the observed dependence on test pulse is due to lack of equilibration at the end of 200 ms test pulse.

Previous studies on Shaker have shown that equilibration of C-type inactivation requires 18 second pulses (Olcese et al., 1997). Although C-type inactivation in Shaker W434F mutant is ultrafast we decided to explore the effects of long test pulse on gating hysteresis. We applied 18 second test pulses from holding potentials of -110 or 0 mV and measured the gating currents upon return to 0 mV (Fig. 3 A). Surprisingly, these currents decay to a stable baseline with little sign of contaminating currents for test pulses in the range of -100 mV to 10 mV (Fig. 3 A). Given the long length of the pulse protocol and the sensitivity of gating current recordings, it is important to check that the maximal gating charge measured using depolarizing and hyperpolarizing pulses is equal. Variable gating charge amplitude could arise if physiological processes occur during the pulse protocol such as endocytosis or changing cellular conditions. Such changes would introduce confounding variables that prevent measurement of Q_SS_-V. Figure S5 shows that the absolute gating charge measured with the return to reference protocol is equal whether measured with depolarizing or hyperpolarizing pulses from a given oocyte despite the long duration of the pulses applied.

**Figure 3:**
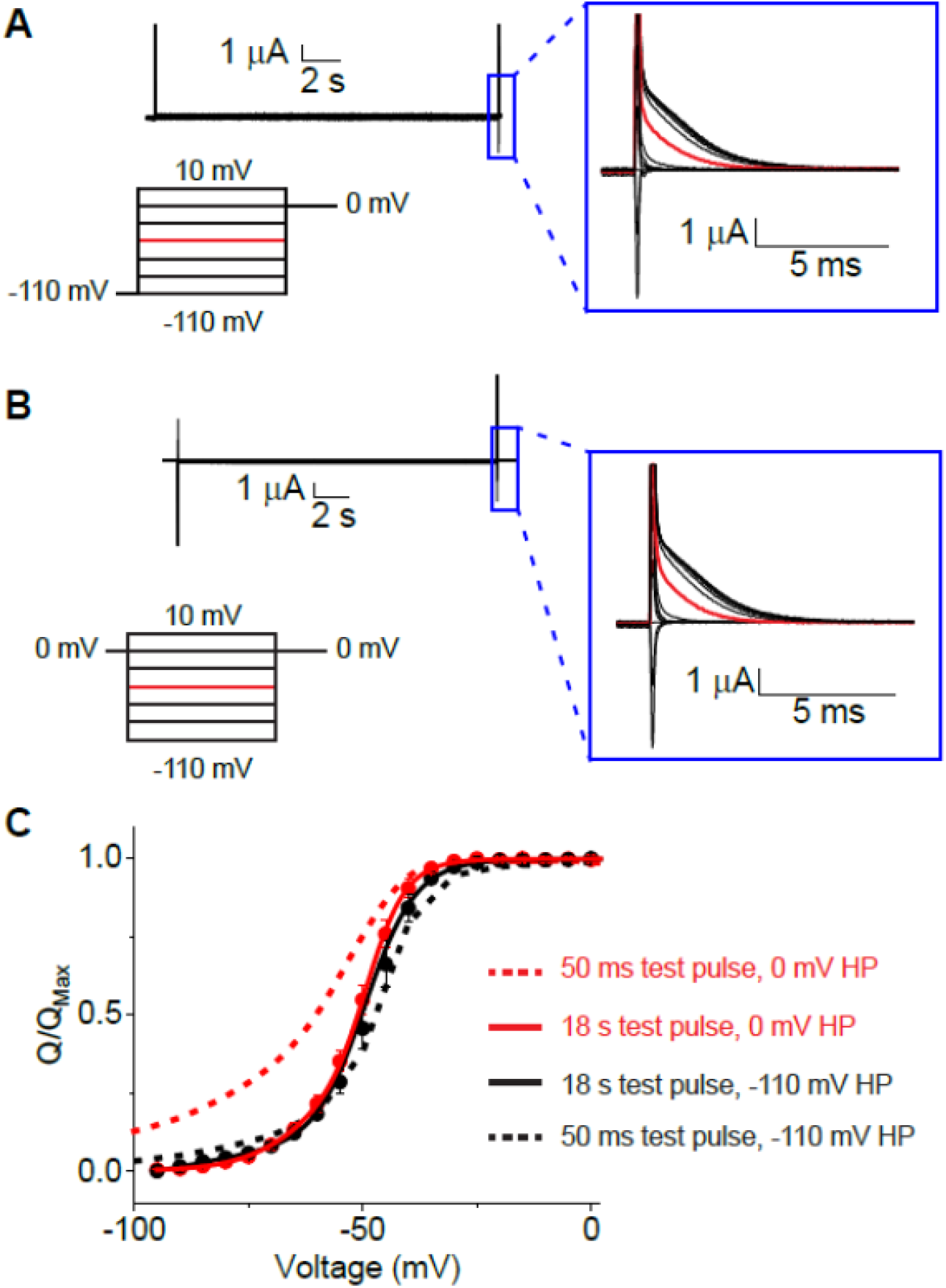
Ultralong pulse duration eliminates hysteresis in Shaker W434F. Raw, unsubtracted recordings of Shaker W434F using 18 second test pulses in the depolarizing (A) or hyperpolarizing (B) direction. Recordings are from the same oocyte and shown on the same scale. In the polarized holding potential condition, a 500 ms prepulse to -110 mV from a holding potential of -80 mV was used to minimize the time the oocyte was held at extreme potentials. Insets at right show an expanded view of the off pulse used for integration of the gating charge. The trace highlighted in red represents the off-gating pulse from the -50 mV test pulse shown on the same scale for comparison. (C) Charge-voltage (Q-V) relationships obtained with 18 second test pulse durations using hyperpolarizing (red) or depolarized (black) test pulses. Dashed lines represent the Q-V curves obtained using 50 ms test pulses for comparison. The difference in V_M_ for the Q-V curves obtained with 18 second test pulses is reduced to 1.8 mV (p = 0.09 in a paired t-test).

The Q-V curves obtained using these 18 second test pulses from depolarized and hyperpolarized potentials nearly overlay (Fig. 3 B). With a separation of just 1.8 mV between the median voltages, the apparent energetic difference observed between activation and deactivation has dropped to 0.5 kcal/mol. The convergence of these curves confirms that hysteresis observed with shorter pulse durations is a consequence of non-equilibrium conditions. Thus, the gating of Shaker W434F is completely reversible and the free energy of activation measured using median voltage analysis under equilibrium conditions captures the entirety of the gating transitions.

### Slow gating kinetics at intermediate potentials cause gating hysteresis

While the above protocol greatly improves the quality of equilibrium gating charge measurements, it comes at the expense of losing information on gating kinetics. To further probe the slow processes limiting channel gating, we used voltage clamp fluorometry. Fluorophore labeling of Shaker A359C W434F provides a spectroscopic probe to track conformational changes within the voltage sensor (Mannuzzu et al., 1996). We recorded VCF signals from TMRM-labeled Shaker A359C-W434F with 25 second test pulses in the depolarizing or hyperpolarizing direction (Fig. 4 A). The stable baseline in the raw fluorescence traces show that there is minimal photobleaching from the dim excitation used. These traces show voltage-dependent quenching of fluorescence at depolarized potentials that is characteristic of Shaker gating. Examining just the first 200 ms of the test pulse (Fig. 4 B) shows that the voltage sensor movement has not equilibrated at intermediate potentials, in agreement with the Q-V curve measurements.

**Figure 4:**
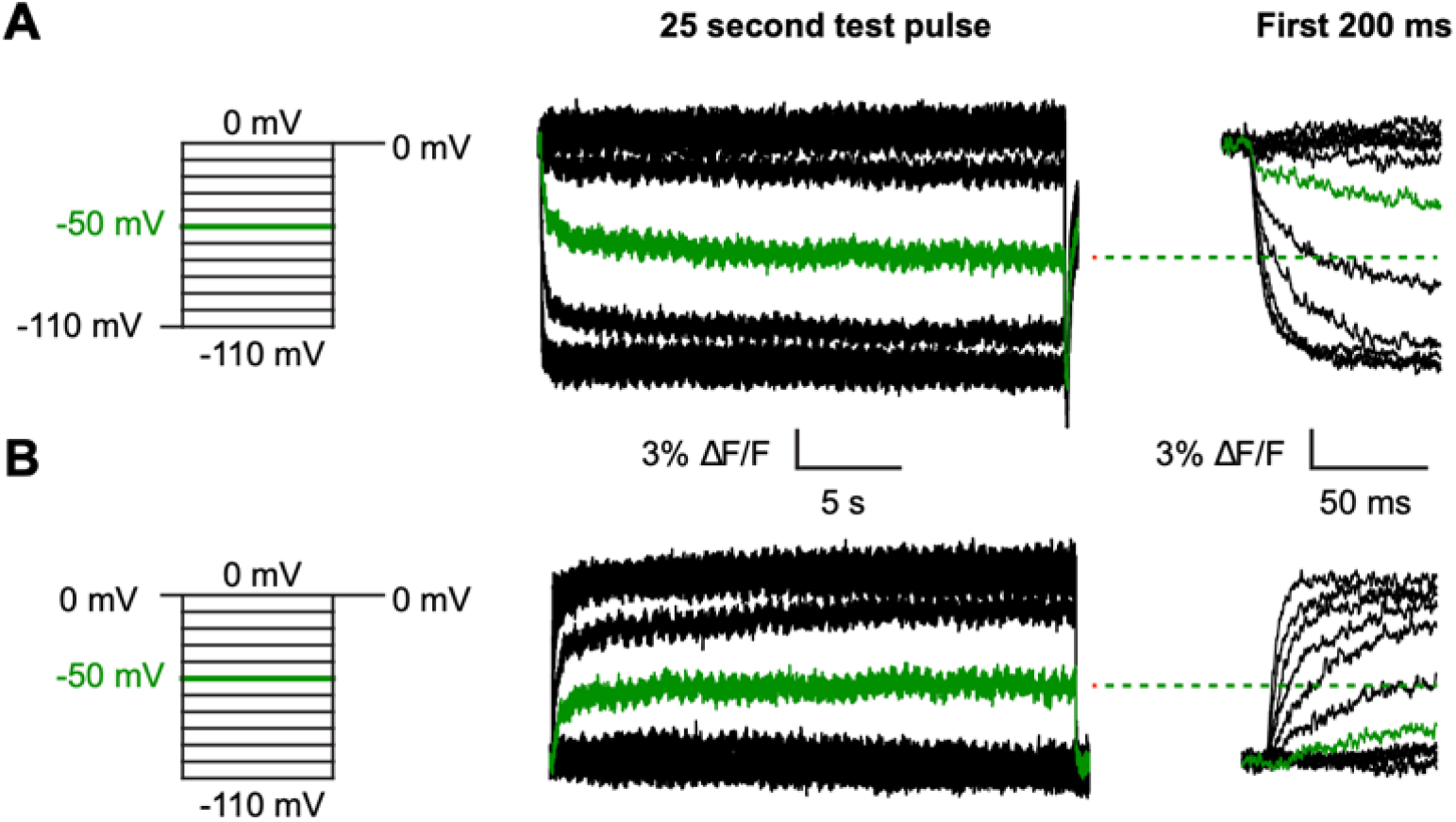
Spectroscopically tracking S4 movement over long pulses VCF protocols (left) and raw fluorescence traces (middle) shown on the linear scale measured with depolarizing test pulses from a potential of -110 mV (A) or hyperpolarizing test pulses from 0 mV (B). The trace highlighted in green in both recordings corresponds to a test pulse of -50 mV. An expanded view of the first 200 ms of the test pulses from A to illustrate the data obtained during typical VCF recordings with dashed lines showing the steady-state position of the red trace at the end of the 25 second pulse.

The normalized fluorescence traces plotted on a log timescale more clearly highlight the slow kinetics limiting equilibration (Fig. 5 A). Test pulses of -40 to -60 mV in either depolarizing or hyperpolarizing direction show fluorescence changes even 10 seconds into the pulse. Despite the slow kinetics of gating at these intermediate potentials, the voltage sensors converge to a similar position whether coming from a depolarized or hyperpolarized holding potential.

**Figure 5:**
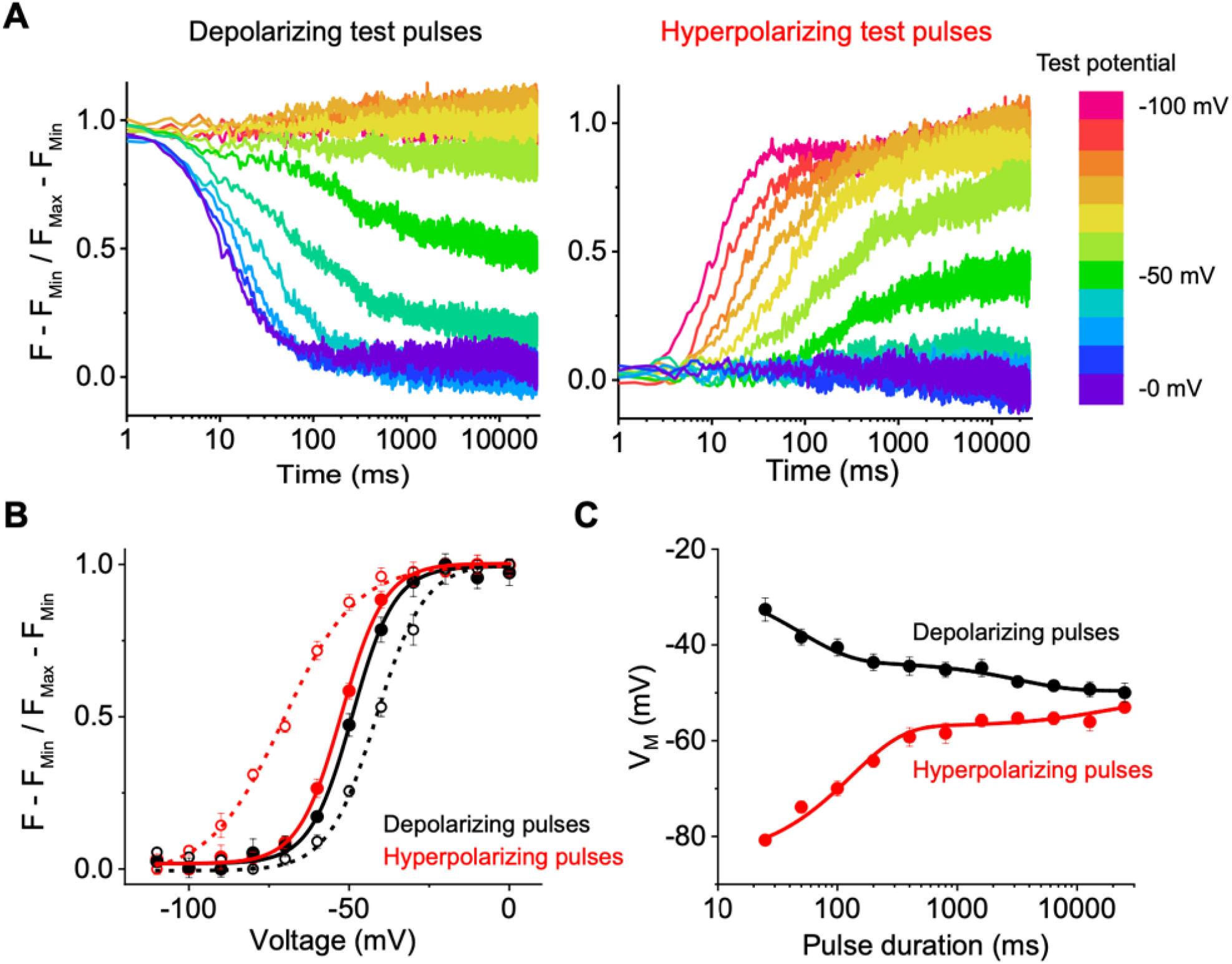
Slow gating kinetics at intermediate potentials underlie hysteresis. (A) Normalized voltage-clamp fluorometry signals for TMRM-labeled Shaker A359C W434F shown on a log timescale for the same oocyte measured with depolarizing (left) or hyperpolarizing (right) test pulses. Test pulses of 25 seconds are color coded according to the scale shown at the far right. The minimal drift in baseline for the traces maintained at the holding potential indicate there is little photobleaching due to the low power of excitation used. F_Min_ corresponds to the average fluorescence of the 0 mV test pulse over the final 100 ms while F_Max_ corresponds to the average fluorescence value of the -100 mv test pulse over the final 100 ms of the pulse. (B) Fluorescence-voltage (F-V) relationships for Shaker A359C W434F extracted at 100 ms (dashed lines) or 25 second (solid lines) pulse duration from polarized (red) or depolarized (black) holding potentials. (C) Median voltage from F-V curves extracted at various time points during a 25 second test pulse from polarized (red) or depolarized (black) holding potentials. Error bars in (B) and (C) show standard error from 3 oocytes. Fits in C represent bi-exponetial decay curves with the following parameters for the black trace: A_1_= 17.1 +/- 3.3, τ_1_= 48.4 +/- 12.8 ms, A_2_= 6.1 +/- 0.8, τ_2_= 3312 +/- 1147 ms, offset= -49.6 +/- 0.6 mV, Χ^2^= 0.22. Parameters for the red trace: A_1_= -28.2 +/- 5.6, τ_1_= 131 +/- 24 ms, A_2_= -5.2 +/- 4.6, τ_2_= 17037 +/- 44714 ms, offset= -51.8 +/- 5.6 mV, Χ^2^= 1.3.

We can monitor the convergence of the system to its equilibrium position by extracting normalized fluorescence-voltage (F-V) curves at various timepoints during the test pulse (Fig. 5 B and 5 C). The 30 mV separation in the F-V curves obtained in the depolarizing versus hyperpolarizing direction at 100 ms is nearly eliminated at the end of the 25 second pulse. This timescale agrees with the elimination of hysteresis in the Q-V curves observed above, indicating that the equilibrium position of the voltage sensor is independent of the holding potential.

The difference in median voltage measured in the depolarizing and hyperpolarizing direction provides a means for comparing the results from gating current and VCF recordings. Although the precise magnitude of the measured hysteresis is different for the two techniques, the timescale for the dissipation is remarkably similar (Fig. 6). The difference in the reported magnitude of hysteresis is not surprising, as the relative fluorescence change between two states does not necessarily correlate with the fraction of charge moved. The time it takes for the voltage sensor movement to equilibrate, however, should be less dependent on the method of detection. The slightly slower convergence of the curves obtained using VCF may reflect slower kinetics for the TMR labeled A359C mutant.

**Figure 6:**
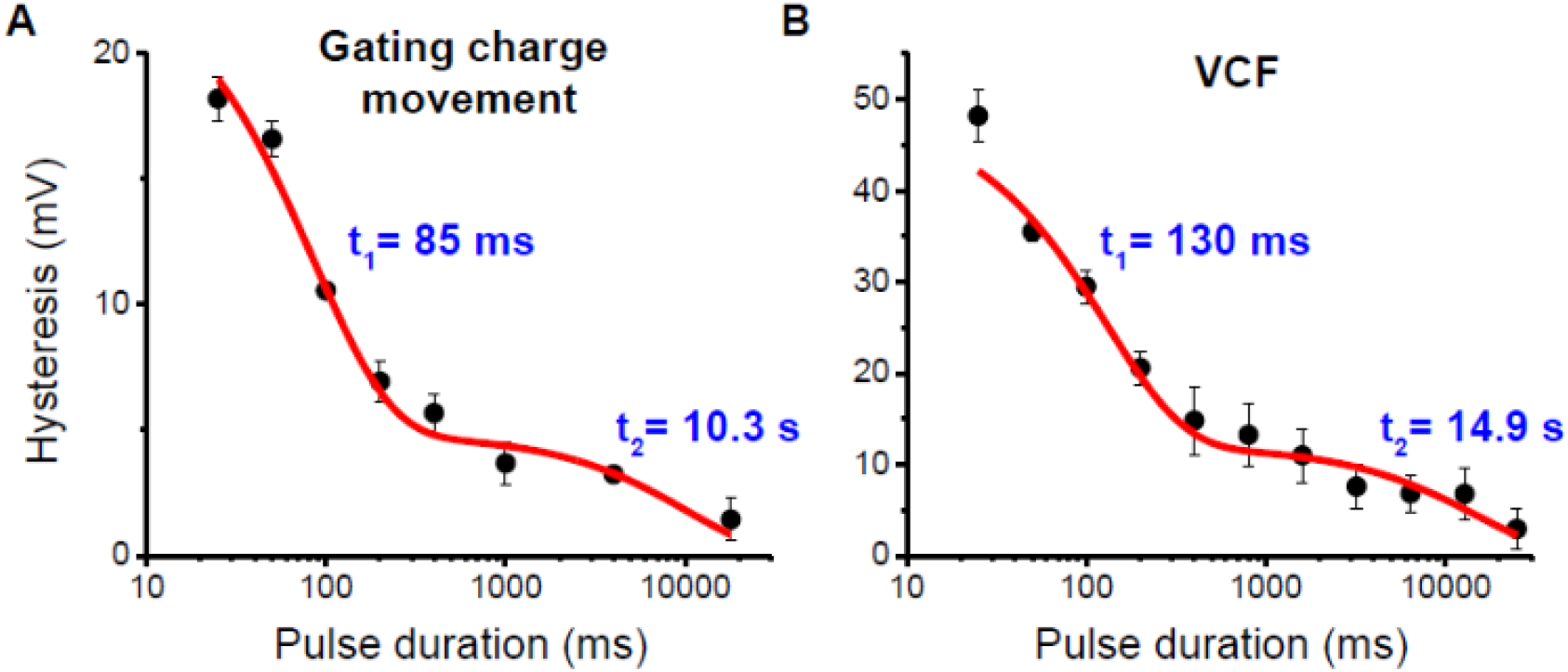
Hysteresis arises due to measurement at non-equilibrium conditions. Hysteresis in (A) charge movement and (B) VCF signal plotted against test pulse duration. Lines represent fits to two-component exponential decays with time constants shown in plots. Parameters for fit in A: A_1_= 19.0 +/- 1.8, τ_1_= 85.0 +/- 11.8 ms, A_2_= 4.8 +/- 0.8, τ_2_= 10305 +/- 4897 ms, Χ^2^= 1.7. Parameters for the fit in B: A_1_= 36.6 +/- 3.3, τ_1_= 129.6 +/- 28.8 ms, A_2_= 12.0 +/- 2.2, τ_2_= 14929 +/- 7531ms, Χ^2^= 1.2. Offset was fixed to 0 mV in both fits. Error bars show standard error from 3 oocytes.

Nevertheless, the agreement between these two techniques strongly suggests that the source of gating hysteresis in Shaker is from non-equilibrium measurements. Approximating equilibrium requires pulse durations far longer than those conventionally used for this channel.

## Discussion

Hysteretic behavior of channel function has a clear physiological significance as it confers “memory” to channel’s gating properties. For channels like hERG (Shi et al., 2020), HCN (Mannikko et al., 2005), and K_v_3.1 (Labro et al., 2015), hysteresis tunes the voltage response of the channel during the course of an action potential waveform in specialized cells and is central to their physiological function. Simulations of SA node activity show that cardiac pacemaking becomes arrhythmogenic if the hysteretic behavior of HCN channels is eliminated or significantly reduced (Mannikko et al., 2005). Hysteresis in channel activity is not surprising because channel activity is a path function and, in most cases, the open channel conformation is not the final state. However, for many voltage-gated ion channels, the net free energy for channel activation appears to be larger than the net free energy of deactivation as the measured Q-V curve shifts depending on the direction the pulses are applied (Haddad and Blunck, 2011; Villalba-Galea, 2017; Villalba-Galea and Chiem, 2020). This difference in free energies is quite surprising because none of the existing gating models (Bezanilla et al., 1982; Schoppa and Sigworth, 1998; Zagotta et al., 1994) can recapitulate hysteresis under equilibrium conditions (Figure S2 and Figure S3 and Appendix). Our studies here show that the hysteresis in Shaker channel gating is due to the inability of the standard gating current protocols to measure the slow charge movements especially at intermediate potentials.

By measuring the charge movement at a suitable reference potential where the ionic currents contamination is minimized and the kinetics of charge movement is relatively rapid, we are able to measure extremely slow charge movement. In a standard gating current protocol, it becomes difficult to measure charge movements that are in the ballpark of hundreds of milliseconds but using a modified protocol, we are able to measure charge movement that occurs in few tens of seconds ^2^.

The choice of the reference potential for measuring gating currents is crucial. In principle, this protocol can be used to measure extremely slow charge movement at any potential as long as the gating currents at the reference potential are fast enough. This is typically not an issue for highly voltage-dependent channels but for a weakly voltage-dependent channel, there is a chance that this optimal reference potential is outside the experimental range.

Our findings that the Shaker Q-V curves do not exhibit hysteresis under equilibrium conditions has number of implications. First of all, our measurements demonstrate that the Shaker gating is not accompanied by any additional dissipation or gain of energy. Recently, it has been argued that the rapid voltage jumps used to estimate charge movement corresponds to non-equilibrium work and, therefore, must be accompanied by dissipation of energy during this process (Villalba-Galea, 2017; Villalba-Galea and Chiem, 2020). According to Villalba-Galea (2017), “as the movement of charges takes places within a condensed system (proteins + water + lipids + ions), friction and non-voltage dependent transitions are expected to dissipate part of the energy gained from the electric work.

Thus, it can be said that

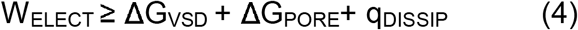

where q_DISSIP_ accounts for the energy dissipated during this process. The inequality highlights that the VSD and surroundings are condensed systems, therefore a fraction of the work is inevitably lost to friction.”

This notion that friction and other non-voltage-dependent transitions cause energy dissipation and,consequently, the net free-energy of activation cannot be measured from the work function for a microscopic process contradicts earlier work by us (Chowdhury and Chanda, 2012) and others (Sigg 2013). If activation of the channel by a rapid voltage pulse causes energy dissipation, then the difference between Q_ON_ and Q_OFF_ reflects a fundamental behavior of the system. However, our demonstration that there exists one and only Q-V curve for a channel irrespective of the holding potential clearly refutes this idea. The charge displaced between two equilibrium ensembles of the system, at two different voltages, will always inform the reversible limit of the total work done associated with the transformation. Energy dissipation or absorption will depend on the path taken by the system but will not be captured by this reversible work. Effects of forces such as friction, solvent viscosity etc. are confined to the kinetics of gating currents rather than equilibrium charge displacement. Thus, regardless of the speed of voltage change, we should be able to determine the free-energy difference between the two states by integrating the gating charge moved at each potential as long as the gating charges settle into their equilibrium position.

Second, our study also shows that the measured steady state Q-V curves are not exclusive reporters of voltage-sensor movement but are also influenced by downstream voltage-independent transitions. Numerous voltage-independent processes such as pore opening (Haddad and Blunck, 2011), inactivation (Olcese et al., 1997), or voltage sensor relaxation (Villalba-Galea et al., 2008) have been proposed as a mechanism for hysteresis in Q-V curves on the basis that these transitions are not observed in gating current recordings. This effect of the voltage-independent transitions can be illustrated by using simple kinetic models where the addition of any voltage-independent transition will induce a shift in Q-V curve even if the microscopic parameters associated with voltage-dependent transition remains unchanged (Fig. S1). While the contribution of these processes to the gating charge movement may be difficult to detect, our data show that using an appropriate recording protocol it may be possible to measure charge movements associated with these processes. These findings also demonstrate that gating charge movement can serve as a reaction coordinate for the whole activation pathway (initiated by voltage). In practice, however, our ability to measure charge movement may be limited by kinetics and signal to noise in the recordings.

Third, our studies establish that at least for the Shaker potassium channel, the observed hysteresis in the Q-V curves stems from the inability of existing measurements to effectively track the slow charge movements during OFF gating. Especially at intermediate potentials where the OFF charge movement is very slow, we find that pulse lengths of ten seconds or more are needed to fully equilibrate the channel. At these potentials, the charge moved is undercounted compared to more extreme potentials where the gating currents become faster. The Q-V curve obtained from depolarized holding potentials, consequently, appears shifted relative to the one obtained from hyperpolarized holding potential. Whether we use the “return to reference” protocol or the standard gating current protocol, the Q-V obtained from hyperpolarized holding potential remains the same.

This is important because the free-energy calculations and gating energetics typically reported in the literature are based on the Q-V curves obtained from hyperpolarized holding potentials (Ryu and Yellen, 2012; Fernandez-Marino et. al. 2018). Therefore, at least for the Shaker K+ channel, the Q-V curves obtained by hyperpolarized holding potentials is similar to the true steady state Q-V curve even when obtained using standard gating current protocol.

It is somewhat surprising that for the Shaker channel, test pulses of about 20 seconds are necessary to equilibrate the channel at intermediate potentials when returning from depolarized potentials. This indicates the channel must traverse through a high energy barrier to return the voltage-sensor to down position. Nonetheless, our ability to measure very slow charge movements using the return to reference protocol suggests that it may be possible to use this protocol to track slow charge movements in other voltage-dependent ion channels. Activity of many members of the VGIC superfamily is highly voltage-dependent but it has not been possible to measure associated gating currents in all cases. In addition, many channels that lack voltage-sensing domain such as K2P channels also exhibit strongly rectifying currents (Schewe et al., 2016) but charge movement associated with channel gating has not been detected in this channel family. The return to reference protocol may allow us to record gating currents from a broader class of channels which are activated by voltage in order to identify the molecular determinants of voltage-sensitivity and shed light on diverse mechanisms of voltage-gating.

## Appendix

Here we use a simple kinetic model to demonstrate that a Q-V curve is not purely a reporter of voltage-dependent transitions but is also influenced by voltage-independent transitions along the activation pathway. This three-state model consists of an initial voltage-dependent transition followed by a voltage-independent transition (Fig. S1).

When a voltage-independent transition is added beyond the open state (such as a relaxed state), the Q-V curve shifts to the left, indicating channel activation becomes more favorable. The magnitude of this shift increases as the favorability of the “relaxed” state relative to the open state is increased. Despite the fact that the voltage-dependent transitions remain unchanged in all the models shown in Fig. S1, the Q-V curves vary widely depending on the properties of the voltage-independent transition. It is important to note that this transition can represent any process, including pore opening, inactivation, ligand binding, or voltage sensor relaxation. All of these processes will influence the Q-V curve if they specifically stabilize one charged state of the channel over another even if they do not carry any charge themselves. This is also apparent if one examines the solution of the differential equations describing the charge movement which contains the eigenvalues of all the transitions including the voltage-independent transitions.

If voltage-independent transitions influence Q-V curves and alter the measured free energy of channel gating, then they cannot be a source of Q-V curve hysteresis. To this point, no balanced kinetic model can produce hysteresis at equilibrium because one of the assumptions inherent to kinetic modeling is that all transitions are microscopically reversible. If all individual transitions are microscopically reversible, then the process as a whole must be thermodynamically reversible and the gating charge of the channel becomes a state function. As a result, even the Q-V curves in a channel with a mode-shifted kinetic regime converge to a single equilibrium position regardless of the direction measured (Fig. S2). Hysteresis in this (or any other) model can only be produced by truncating the gating current recording before the charge movement has equilibrated at all potentials.

Inversely, non-equilibrium measurements result in Q-V curve hysteresis regardless of the model, even for a two-state model (Fig. S3). The Q-V curve hysteresis obtained from a single non-equilibrium test pulse duration for a two-state model is difficult to distinguish from hysteresis observed in a mode-shifted gating model (comparing Fig. S2 B and Fig. S3 B). Thus, Q-V curve hysteresis observed using a single test pulse duration provides little to no information about the channel gating model. Examining the time course of disappearance of hysteresis at intermediate potentials, however, provides information of the slow processes limiting equilibrium that can better inform kinetic modeling efforts (comparing Fig. S2 C and Fig. S3 C).

An interesting phenomenon observed in our experimentally measured Q_ss_-V curves is that the curve measured with depolarizing pulses displays faster apparent convergence to the equilibrium value than the curve measured with hyperpolarizing pulses. The simplest mechanism that can account for this phenomenon is an asymmetry in the charge moved to the transition state in the forward and reverse directions, as this can produce asymmetry in convergence to equilibrium even in a two-state model (Fig S4). This results in differing voltage dependencies in the rates of the forward and reverse transitions, and if the ttransition state is closer in charge to the open state than closed, the Q-V curves measured in the forward direction (depolarizing pulses) will be a better approximation for the QEQ-V curve than the reverse (Fig S4). More complex models such as mode-shift or VSD-relaxation schemes can also account for this phenomenon, but this two-state model is the simplest mechanism that can account for such an asymmetric convergence to equilibrium.

## Acknowledgements

We are grateful to Chris Lingle, Francisco Bezanilla, Sandipan Chowdhury, Gail Robertson, Lucie Delemotte and members of the Chanda lab for their helpful feedback at various stages. This work was supported by funding from the National Institutes of Health (NS101723, NS081293, NS116850) to B. Chanda and J. Cowgill (T32 HL-07936-17).

## Supplemental figures

**Figure S1:**
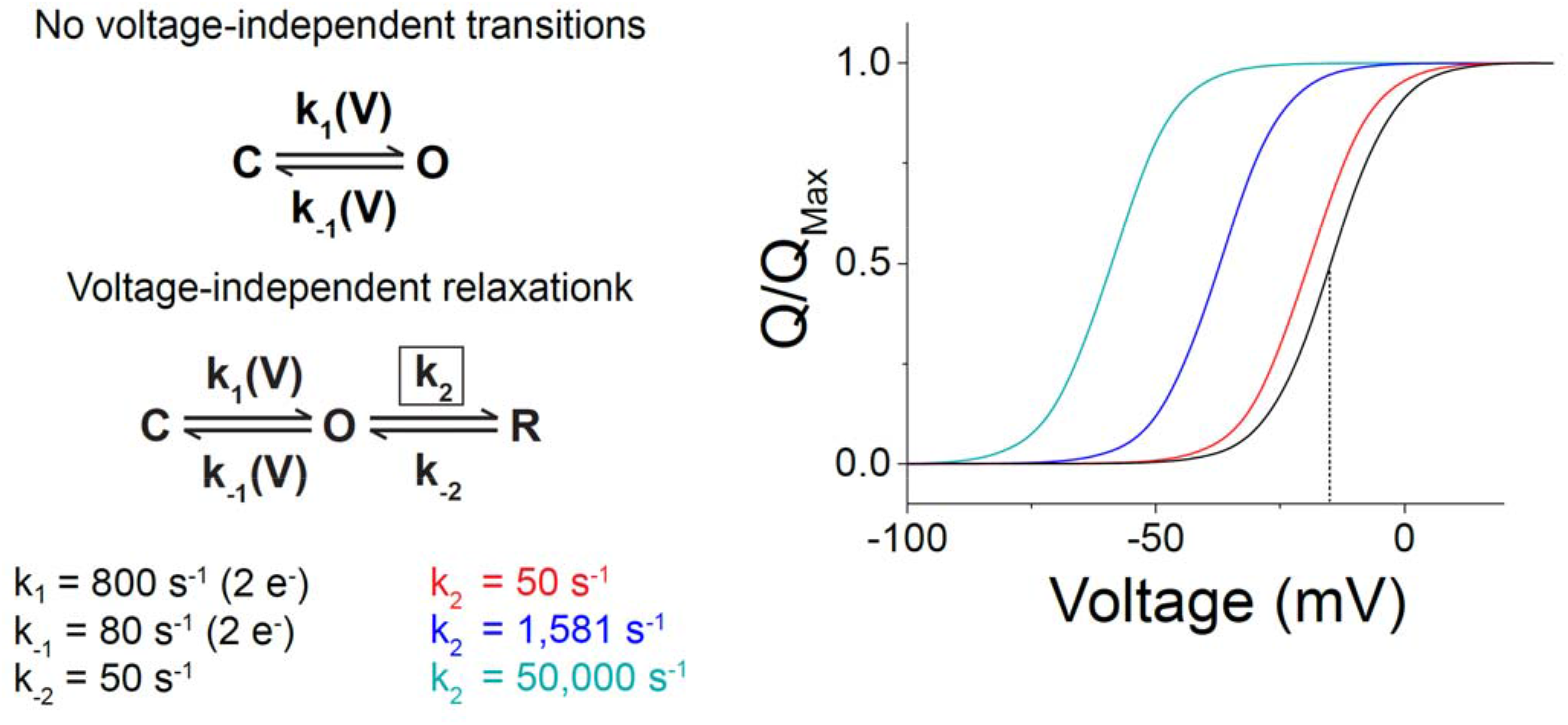
Voltage-independent processes influence Q-V curves Left: Models and rate constants used in simulation of steady-state Q-V curves at right. Note that the voltage-dependent transitions are the same in all models and only voltage-independent rates vary. Right: Simulated Q-V curves for the two-state (black) and three-state (colors) models in part A. The blue trace corresponds to a ΔG_relaxation_ of -2 kcal/mol, the cyan trace corresponds to ΔG_relaxation_ of -4 kcal/mol, and the red trace corresponds to ΔG_relaxation_ of 0 kcal/mol. Despite the lack of charge movement associated with the relaxed state, it still influences the Q-V curve because it preferentially stabilizes the activated voltage sensor.

**Figure S2:**
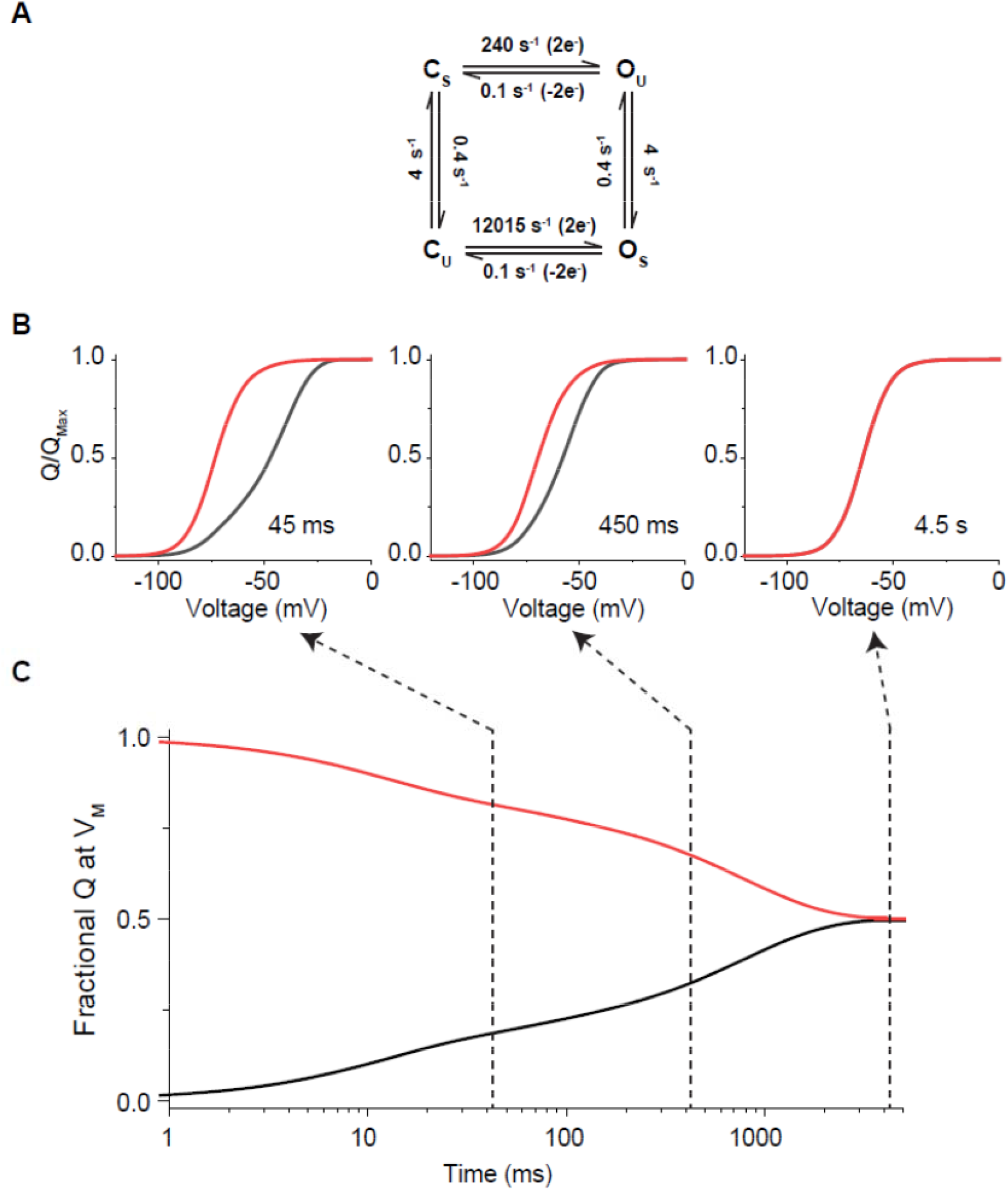
Mode shift does not generate hysteresis in Q-V curve at equilibrium. (A) Four-state kinetic model representing a classical mode-shifted gating scheme used for simulating data in parts B and C according to similar protocol as Figure 2A. (B) Q-V curves extracted at various time points during the simulated test pulses in A from a holding potential of -100 mV (black trace) or 0 mV (red trace). (C). Time course of the integrated gating charge movement for the pulse to -64 mV (V_M_ of the channel) from a holding potential of -100 mV (black trace) or 0 mV (red trace). Dashed lines indicate the points where the Q-V curves in part B are extracted.

**Figure S3:**
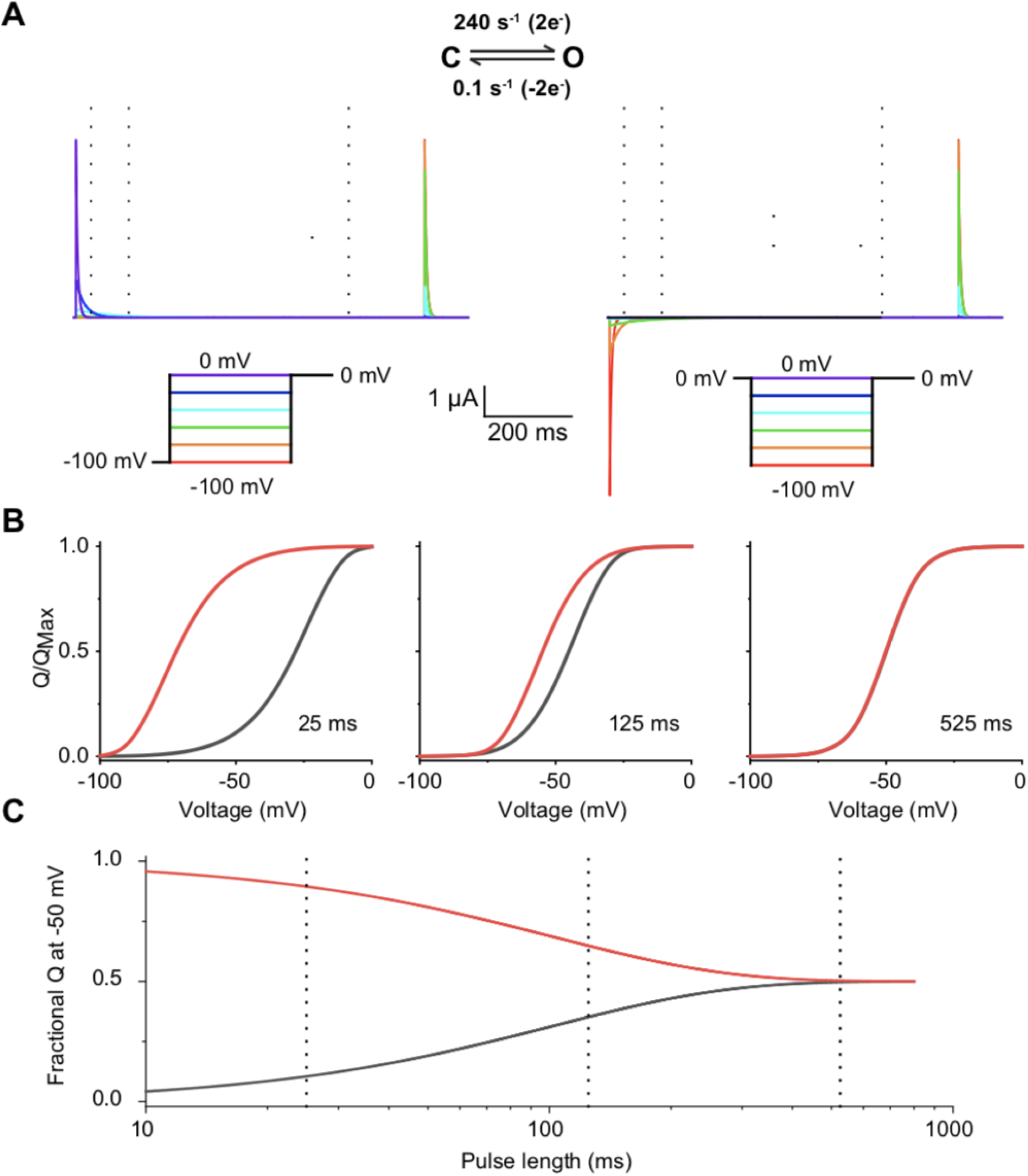
Two-state models can also produce Q-V curve hysteresis under non-equilibrium conditions. Two-state kinetic model (top) and gating currents simulated according to the inset protocols (bottom). Dotted lines indicate the length of the pulse integrated to produce the Q-V curves in part B. Q-V curves extracted at various time points during the simulated test pulses in A from a holding potential of -100 mV (black trace) or 0 mV (red trace). (C). Time course of the integrated gating charge movement for the pulse to -50 mV from a holding potential of -100 mV (black trace) or 0 mV (red trace). Dashed lines indicate the points where the Q-V curves in part B are extracted.

**Figure S4:**
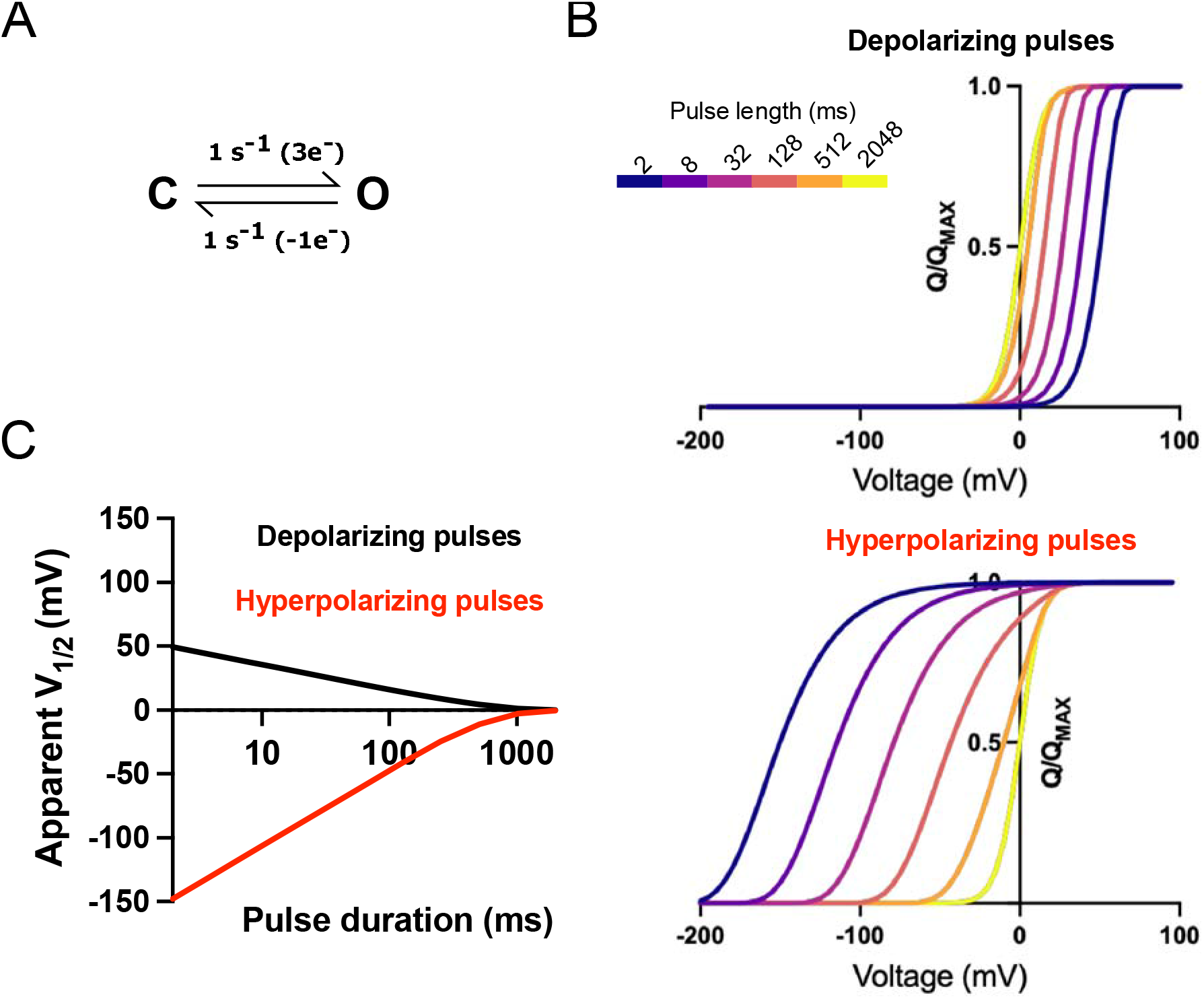
Two-state model can account for asymmetric convergence to equilibrium (A) Two-state model for channel gating with an asymmetric transition state that is closer in charge to the open state (O) than closed (C), as shown by the 3 e-voltage-dependence to the forward rate compared to only 1 e-voltage-dependence to the reverse. (B) Q-V curves simulated with depolarizing pulses from negative holding potentials (top) and hyperpolarizing pulses from positive potentials (bottom) colored according to the pulse duration based on the inset color bar. (C) Apparent V1/2 for the simulated Q-V curves as a function of test pulse duration for model in A simulated with depoalrizing (black) or hyperpolarizing (red) test pulses.

**Figure S5:**
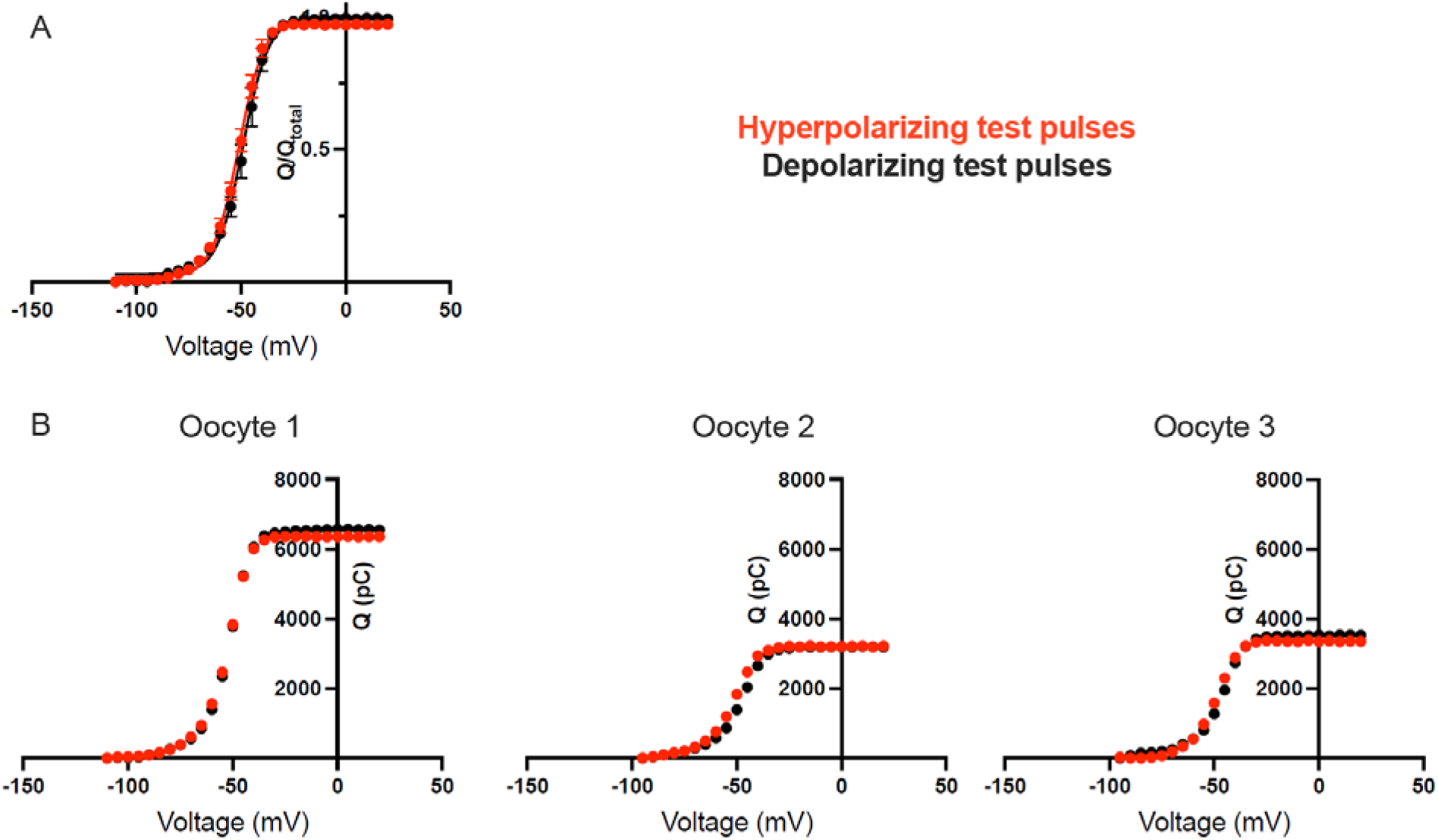
Q-V curves measured from hyperpolarizing and depolarizing test pulses using return to reference protocol. (A) Normalized Q-V curves from hyperpolarized and depolarized holding potentials using return to reference protocol from three independent measurements. (B) Each panel shows Q-V curve from a single oocyte for hyperpolarized and depolarized holding potentials. Over the course of the measurement, the total charge on the membrane does not change.

